# Temperature-dependent intracellular crystallization of firefly luciferase in mammalian cells is suppressed by D-luciferin and stabilizing inhibitors

**DOI:** 10.1101/2024.05.06.592811

**Authors:** Haruki Hasegawa

## Abstract

Firefly luciferase (Fluc) from *Photinus pyralis* is one of the most widely used reporter proteins in biomedical research. Despite its widespread use, Fluc’s protein phase transition behaviors and phase separation characteristics have not received much attention. Current research uncovers Fluc’s intrinsic property to phase separate in mammalian cells upon a simple cell culture temperature change. Specifically, Fluc spontaneously produced needle-shaped crystal-like inclusion bodies upon temperature shift to the hypothermic temperatures ranging from 25°C to 31°C. The crystal-like inclusion bodies were not associated with or surrounded by membranous organelles and were likely built from the cytosolic pool of Fluc. Furthermore, the crystal-like inclusion formation was suppressed when cells were cultured in the presence of D-luciferin and its synthetic analog, as well as the benzothiazole family of so-called stabilizing inhibitors. These two classes of compounds inhibited intracellular Fluc crystallization by different modes of action as they had contrasting effects on steady-state luciferase protein accumulation levels. This study suggests that, under substrate insufficient conditions, the excess Fluc phase separates into a crystal-like state that can modulate intracellular soluble enzyme availability and protein turnover rate.

## 1. Introduction

Luciferase (EC 1.13.12.7) from the North American firefly, *Photinus pyralis*, is a widely used reporter protein that has established its value as an indispensable molecular tool for life science, biomedical research, and drug discovery screening assays [1–4]. Because of its importance in modern scientific research, various aspects of firefly luciferase protein have been extensively characterized [4–7]. Several important features make firefly luciferase a preferred reporter system for biomedical applications. The first and foremost is its high assay sensitivity. Unlike fluorescent reporter systems that can be hindered by the intrinsic autofluorescence of biological materials, mammalian cells and model animals are usually devoid of endogenous bioluminescence; and this ensures an excellent signal-to-background ratio [2]. Another important aspect of firefly luciferase is its short intracellular half-life (∼3 hr [8]), which makes it a reliable reporter that reflects timely transcriptional activity changes regulated by the response elements activated by cognate stimuli. In addition, various luciferase mutant variants have been engineered [2, 9], and different synthetic substrates were produced [2, 10, 11] to enhance this platform’s versatility.

In nature, firefly luciferase is a peroxisome-localizing protein expressed in the photocytes of the firefly lantern organ [12, 13]. The findings that recombinant firefly luciferase can be targeted correctly to peroxisomes in heterologous mammalian cell lines led to the identification of the C-terminal SKL tripeptides as the highly conserved Type-1 peroxisome targeting signal (PTS1) [14, 15]. Because PTS1 is located at the C-terminus, the PTS1-mediated peroxisome import is a post-translational event that comes into effect after the luciferase polypeptide is fully synthesized in the cytosol [16–18]. PEX5, the cargo receptor for PTS1-dependent peroxisome import, then captures PTS1-containing cargo in the cytosol and shuttles it to the docking complex on the peroxisome membrane [19]. Recombinant overexpression studies in mammalian cells revealed that this PTS1 transport pathway is easily saturable [20, 21] as the excess luciferase overwhelms the PEX5 capacity and accumulates in the cytosol (instead of efficiently sorted into peroxisomes) and masking the peroxisome-localizing counterpart when expressed at a very high level [22].

Despite the increasing use of firefly luciferase reporters in numerous applications, the details of luciferase protein phase behaviors and phase separation characteristics have not received much attention except in a few studies. In 1956, by using the native firefly luciferase purified from 30 grams of dried lantern organs isolated from 6,000 fireflies, Green and McElroy reported that purified firefly luciferase readily produced needle-shaped crystals when dialyzed against a low ionic neutral pH buffer [23]. More recently, in 2015 and 2020, Redecke’s group reported that recombinant firefly luciferase spontaneously crystallized intracellularly when overexpressed in baculovirus-infected insect cell lines Sf9 [24] and High-Five [25], in which luciferases produced rod-shaped crystalline inclusions that sometimes reach >100 μm in length. While these observations suggested that firefly luciferase might have an intrinsic propensity to condense or phase separate readily into protein crystals under certain conditions *in vitro* and *in cellulo*, it also posed an interesting question: why such phenomena have been unrecognized in more widely used mammalian cells or transgenic animals. Whether this intracellular luciferase crystallization event requires insect-specific co-factors or insect-like cellular environments is currently unknown. One possibility is that unidentified insect factors that support luciferase crystallization are missing in mammalian cells. Conversely, inhibitory factors are present in mammalian cells and prevent luciferase from crystallizing *in cellulo*. An equally likely possibility is that crystallizing propensity is embedded in the luciferase protein itself, but such propensity appears only under specific cellular environments or growth conditions.

This study aims to elucidate the cellular requirements that support and modulate firefly luciferase crystallization using a mammalian cell host. To this end, firefly luciferase is recombinantly overexpressed in HEK293 cells which previously facilitated the intracellular crystallization of various model proteins [26–31]. The transfected cells are simply cultured at four different settings: (1) regular mammalian cell growth conditions at 37°C with 5% CO2, (2) mild hypothermic condition at 31°C with 5% CO2, (3) 27°C with ambient CO2, and (4) 25°C with 5% CO2 using a cooling-capable incubator. When overexpressed, luciferase mostly distributes to the cytosol regardless of the cell culture temperatures. At 37°C, luciferase produces aggresome-like globular cytosolic inclusion bodies in roughly half of the transfected cell population.

Conversely, at hypothermic temperatures of 31°C and below, luciferase generates needle-shaped crystal-like inclusion bodies that often exceed 100 μm in length. The needle-like inclusion bodies are not associated with or surrounded by membranous organelles and are thus likely built from the cytosolic pool of luciferase. Furthermore, the crystal-like inclusion body formation at hypothermic temperatures is effectively prevented by culturing the transfected cells in the presence of D-luciferin substrate and its synthetic analog, as well as the benzothiazole family of so-called stabilizing inhibitors. Interestingly, D-luciferin and stabilizing inhibitors show substantially different effects on luciferase steady-state intracellular accumulation, thereby suggesting their different modes of action in suppressing luciferase crystallogenesis. These findings unexpectedly identify luciferase’s novel temperature-dependent liquid-to-solid phase separation propensity. Given that the crystal-like phase transition state is only triggered in the absence of substrate, firefly luciferase can potentially use this switch-like mechanism to control intracellular enzyme availability by regulating its solubility and half-life during substrate insufficiency. Additionally, this study demonstrates mammalian cells as a versatile “test tube” to accumulate recombinant intracellular proteins by diverting the cargo protein of interest from the normal protein turnover cycle into a more stable storage pathway in the form of protein crystals.

## 2. Results

### 2.1. Overexpressed firefly luciferase accumulates in the cytosol of HEK293 cells as aggresome-like inclusion bodies under normal cell culture conditions

To study the intracellular phase behavior of firefly luciferase (EC 1.13.12.7) from a North American firefly *Photinus pyralis*, I used the pTT^®^5 vector and HEK293-6E cell overexpression platform (see Materials and Methods). This robust recombinant protein expression system was used previously to induce intracellular crystallization of human Charcot-Leyden Crystal (CLC) protein [30], a cataract-causing R37S mutant of human crystallin-γ-D (CRYGD) [30], and baculovirus polyhedrin [30] in the cytosol and nucleoplasm. Similarly, the same expression system allowed the crystallization of organelle resident proteins and secretory proteins such as human and mouse NEU1 [30], *Anomala cuprea* entomopoxvirus fusolin [31], a human λLC clone [31], and several human IgG monoclonal antibodies [26, 27, 29] in the ER lumen.

When expressed using this platform, a full-length firefly luciferase (∼62 kDa) was readily detectable in cell lysates by SDS-PAGE (Fig. 1A). The quantity of luciferase was equivalent to one of the most abundant endogenous proteins, Hsp70 (Fig. 1A, see the band just above luciferase). At this expression level, firefly luciferase showed cytosolic distribution (Fig. 1B) instead of a peroxisome-like dispersed punctate staining pattern (see below). One likely explanation for the cytosolic accumulation is that the post-translational cytosol-to-peroxisome protein import mechanism via the peroxisomal targeting signal-1 (PTS1) pathway [14, 16] was overwhelmed due to abundant luciferase synthesis in the cytosol. As a result, the majority of luciferase remained in the cytosol and masked the peroxisome-localizing species. In fact, the cytosolic localization of luciferase agreed with an earlier observation in transiently transfected monkey CV-1 cells where the protein expression level influenced the luciferase subcellular distribution between peroxisomes and the cytosol [22]. Of note, in about 50% of transfected HEK293 cells, firefly luciferase formed aggresome-like inclusion bodies readily discernible in DIC images (Fig. 1B, arrowheads). When visualized by PEX14 immunofluorescent staining, peroxisomes showed punctate morphology and dispersed throughout the cytoplasm in both transfected and non-transfected cells (see Fig. 1C). However, when the aggresome-like inclusion body was present, peroxisomes not only clustered near the inclusion body but also spatially overlapped with the globular inclusion (Fig. 1C, arrowheads). The distribution and morphology were also affected in other organelles. For example, the ER membranes (calnexin) and lysosome membranes (LAMP1) became closely juxtaposed to the inclusion body and sometimes encapsulated it (Fig. 1D, arrowheads). Likewise, the *cis*-Golgi membrane (giantin) and *trans*-Golgi membrane (p230) were closely positioned against the inclusion bodies (Fig. 1D).

**Figure 1.**
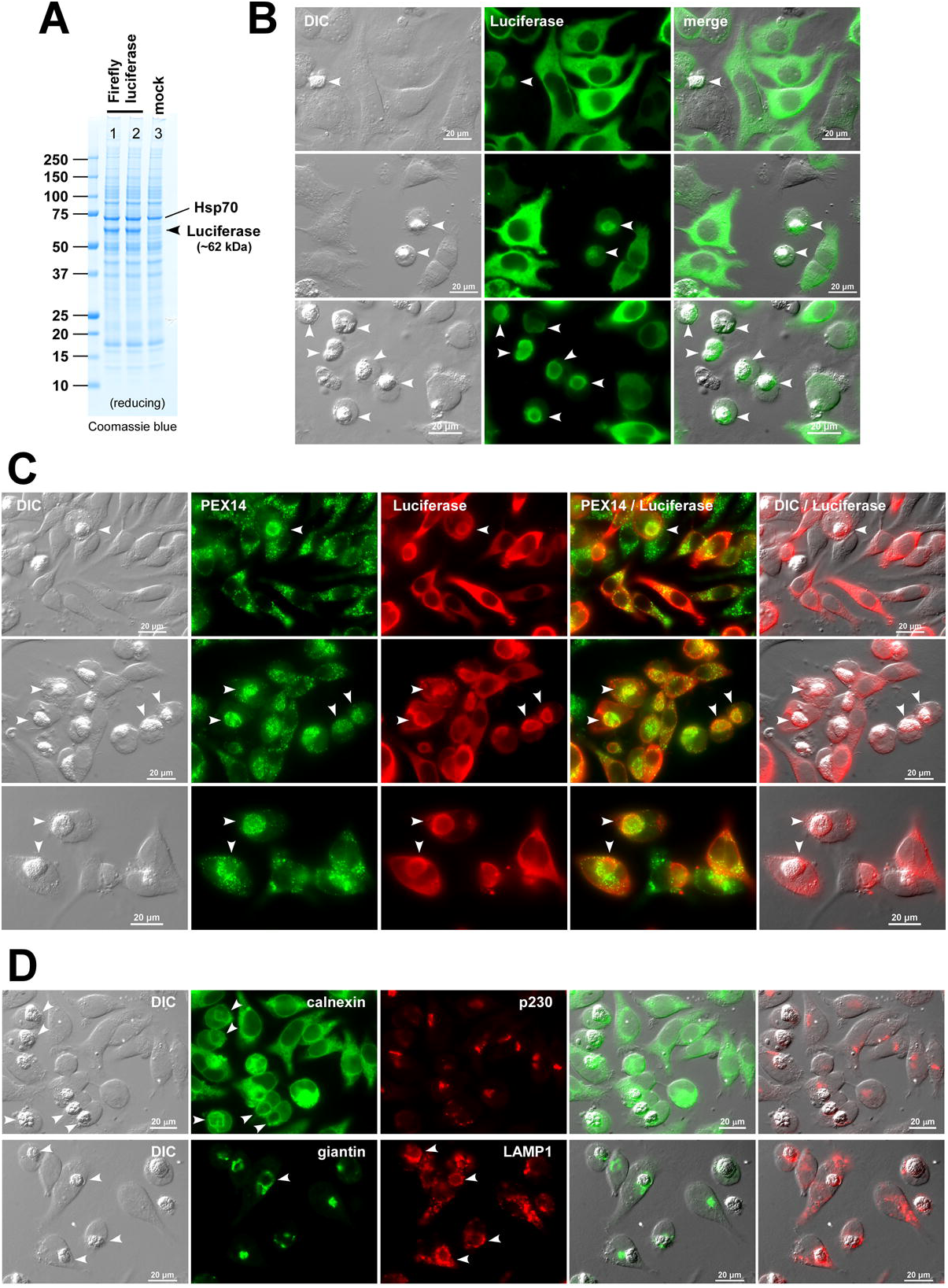
Firefly luciferase accumulates in the cytosol of HEK293 cells. (A) Coomassie blue staining of the whole cell lysate samples prepared from firefly luciferase-expressing cells (lanes 1–2) or mock-transfected cells (lanes 3). Cell lysates were prepared at 72 hours post-transfection. The Firefly luciferase protein band is pointed by an arrowhead. A prominent protein band for endogenously expressed Hsp70 is also indicated. (B) Fluorescent micrographs of HEK293 cells transfected to overexpress firefly luciferase at 37°C cell culture conditions. On day-3 post-transfection, cells were fixed, permeabilized, and immunostained with goat anti-luciferase antibody. Aggresome-like prominent inclusion bodies are pointed by arrowheads. DIC and green image fields were digitally overlayed to create ‘merge’ views. (C) Co-staining was performed using rabbit anti-PEX14 antibody (green) and goat anti-luciferase antibody (red). Green and red image fields were digitally merged in the fourth column. DIC and red image fields were merged in the fifth column. Aggresome-like inclusion bodies are pointed by arrowheads. (D) Luciferase overexpressing cells were co-stained with rabbit anti-calnexin (green) and mouse anti-p230 (red) antibodies (top row). Likewise, co-stained with rabbit anti-giantin (green) and mouse anti-LAMP1 (red) antibodies (bottom row). Calnexin, giantin, and LAMP1 membranes that encircle the aggresome-like inclusion bodies are pointed by arrowheads.

### 2.2. The overexpressed firefly luciferase phase separates into needle-shaped inclusion bodies in hypothermic cell culture conditions

Based on the precedents in insect cell recombinant expression [24, 25], I naively assumed that firefly luciferase would readily crystallize into needle-shaped inclusion bodies by simply overexpressing it in mammalian cells. However, as shown above, luciferase failed to crystallize in HEK293 cells even at a high enough intracellular concentration at which half of the transfected cell population developed aggresome-like inclusion bodies. In other words, a high enough luciferase protein concentration alone was not sufficient to drive luciferase crystallization inside the transfected mammalian cells. Because insufficient expression level seemed unlikely to explain the lack of luciferase crystallization in HEK293 cells, I looked for other potential reasons that could account for luciferase’s phase behavior discrepancy between mammalian cells and insect cells.

Besides the cell hosts and the cell culture media, obvious differences between insect cell and mammalian cell culture conditions are the growth temperatures (i.e., 37°C vs 27°C) and the presence or absence of 5% CO2 overlay. When it comes to a gene delivery method, the insect cell system relies on recombinant baculovirus to introduce and express the gene of interest [24, 25], while HEK293 cells rely on plasmids and a polycation-based transfection system [26–31]. To probe if insect cell culture temperature or ambient CO2 level would influence the intracellular phase behavior of firefly luciferase, a shake flask of transfected HEK293 suspension culture was directly transferred from a regular mammalian cell culture incubator (37°C, 5% CO2) to an insect cell culture incubator (27°C, ambient CO2) on day-1 post-transfection. Surprisingly enough, rod-shaped inclusion bodies started forming spontaneously inside the suspension cultured HEK293 cells within 72 hours after shifting to the insect cell growth condition (Fig. 2A). Due to the considerable length of the inclusion body, crystal-housing cells were easily recognizable during a routine cell counting and viability measurement using trypan blue. Furthermore, HEK293 cells remained mostly viable for at least 5 days in this insect cell growth conditions, and the rod-shaped inclusion bodies continued to elongate during this period. Although firefly luciferase crystallization was induced in HEK293 cells by simply growing the cells in insect cell culture conditions, it was still unclear whether the temperature shift from 37°C to 27°C or the change in CO2 concentration from 5% to the ambient level (= 0.04%) served as the cue for luciferase crystallization. Importantly, however, it became certain that luciferase crystallization in HEK293 cells did not require additional extrinsic factors such as insect cell-specific proteins or baculovirus-encoded gene products.

**Figure 2.**
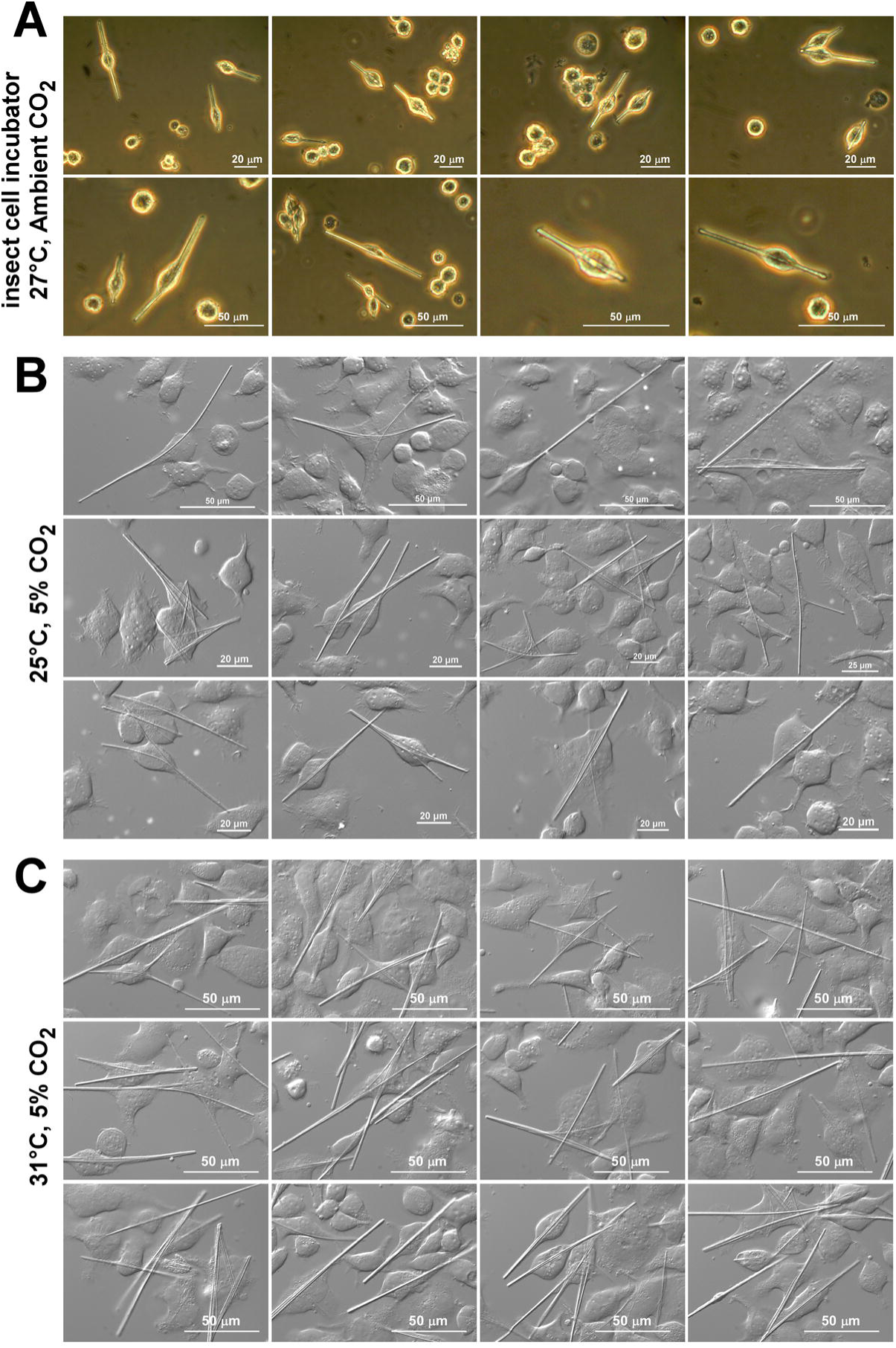
Firefly luciferase phase separates into needle-shaped inclusion bodies in HEK293 cells when incubated at hypothermic growth conditions. (A) At 24 hours post-transfection, a flask of suspension cultured HEK293 cells was transferred to a shaker platform located in an insect cell incubator adjusted to 27°C with ambient CO2. After 72 hours of growth in the insect cell incubator, suspension cultured HEK293 cells were directly viewed using a phase contrast microscope. (B, C) After growing at the regular cell culture condition of 37°C for 24 hours, transfected cells were seeded onto glass coverslips, and transferred to cooling incubators attuned to 25°C or 31°C at 5% CO2. After 72 hours of cell growth at hypothermic 25°C and 31°C temperatures, cells were fixed with paraformaldehyde, and images were captured on day-4 post-transfection using a DIC microscope.

To assess the effect of different cell culture temperatures on luciferase crystallization events while maintaining the constant 5% CO2 overlay, HEK293 cells were first transfected and maintained under the normal cell culture condition (37°C, 5% CO2) for 24 hours before the cells were transferred to a different incubator adjusted to 31°C with 5% CO2 or to a cooling CO2 incubator maintained at 25°C with 5% CO2. At 31°C and 25°C, firefly luciferase induced needle-shaped crystal-like inclusion bodies abundantly in HEK293 cells (Fig. 2 BC). It was, therefore, the cell culture temperature, instead of the low CO2 concentration, that played a role in intracellular luciferase crystallogenesis. In addition, when cell lysates were analyzed by Western blotting up to day-6 post-transfection using three different polyclonal antibodies against firefly luciferase, overexpressed luciferase remained intact (∼62 kDa) without producing any notable degradation products at different cell growth temperatures (data not shown). This precluded the role of degradation products, if any, that might have selectively promoted or inhibited luciferase crystallization at certain cell culture temperatures.

In summary, firefly luciferase did not undergo a crystallization-like phase separation process when luciferase-expressing cells were cultured at a normal cell growth temperature of 37°C; yet needle-like inclusion bodies readily and spontaneously formed when the cells were grown at mild hypothermic conditions such as 25°C, 27°C, or 31°C irrespective of suspension or adherent cell culture format. The morphology of the needle-like inclusion bodies in HEK293 cells was indistinguishable from the needle-like crystals produced by the firefly luciferase overexpressed in Sf9 and High Five insect cells [24, 25]. These needle-like inclusion bodies produced in mammalian cells were also identical to the crystals generated from the native firefly luciferase purified from dried lantern organs isolated from fireflies [23, 32].

### 2.3. Needle-like inclusion bodies are built from the cytosolic pool of luciferase

To assess whether the needle-shaped inclusion bodies are associated with any specific organelle membranes such as the ER or peroxisomes, various subcellular markers were visualized by immunofluorescent microscopy. Similar to the normal cell culture condition at 37°C (see above, Fig. 1), firefly luciferase showed cytosolic distribution at 25°C irrespective of the crystal formation (Fig. 3A). Immunostaining using anti-luciferase antibodies failed to stain the needle-like inclusion body itself (Fig. 3A), which suggested that detection epitopes were lost or masked upon liquid-to-solid phase separation. Similar observations were commonly reported for various intracellular protein crystals [26–31]. Co-staining experiments revealed that inclusion bodies did not show notable associations with the membranes of the ER (calnexin), *cis*-Golgi (giantin), *trans*-Golgi (p230, TGN46), lysosomes (LAMP1), mitochondria (Tom20), and nuclear membrane (Nup153), as well as proteinaceous polymers such as microtubules (β-tubulin) (see Fig. 3B).

**Figure 3.**
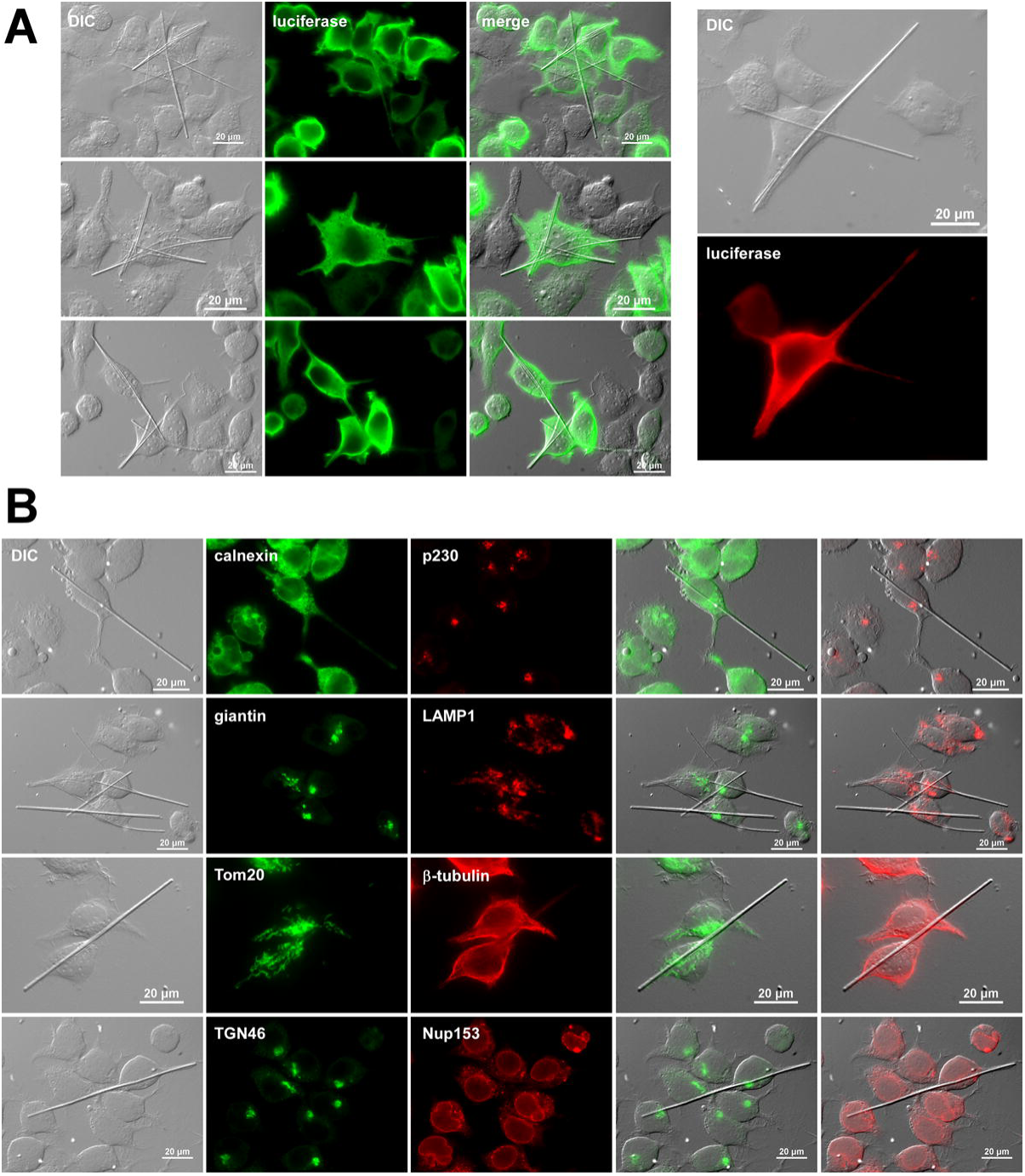
Needle-like inclusion bodies are not associated with organelle membranes or microtubule cytoskeleton. (A, left panel) Transfected cells were seeded onto glass coverslips and transferred from 37°C to 27°C culture condition 24 hours after transfection. After 72 hours at hypothermic cell growth conditions (on day-4 post-transfection), cells were fixed, permeabilized, and immunostained with goat anti-luciferase polyclonal antibody and Alexa Fluor 488 anti-goat secondary antibody. DIC and green image fields were digitally overlayed to create ‘merge’ views. (A, right panel) Transfected cells were stained with rabbit anti-luciferase monoclonal antibody and Alexa Fluor 594 anti-rabbit secondary antibody. (B) Co-staining was performed using rabbit anti-calnexin antibody and mouse anti-p230 antibody (top row). Rabbit anti-giantin antibody and mouse anti-LAMP1 antibody (second row). Rabbit anti-Tom20 and mouse anti-β-tubulin (third row). Rabbit anti-TGN46 and mouse anti-Nup153 (bottom row). DIC and green image fields were digitally merged to create the fourth column. DIC and red image fields were merged to create the fifth column.

Similarly, co-staining with a peroxisome membrane marker PEX14 did not reveal notable associations with the needle-like inclusion bodies (Fig. 4 AB). Because of the punctate nature of peroxisome morphology, it was difficult to postulate that peroxisome membranes can stretch or homotypically fuse to enclose a needle that often reaches 100 μm in length (Fig. 4 AB). Based on its predominant cytosolic localization in conjunction with the apparent lack of association with the organelle membrane system, the crystal-like inclusion bodies were most likely formed in the cytosol using the cytosolic pool of luciferase as the building block.

**Figure 4.**
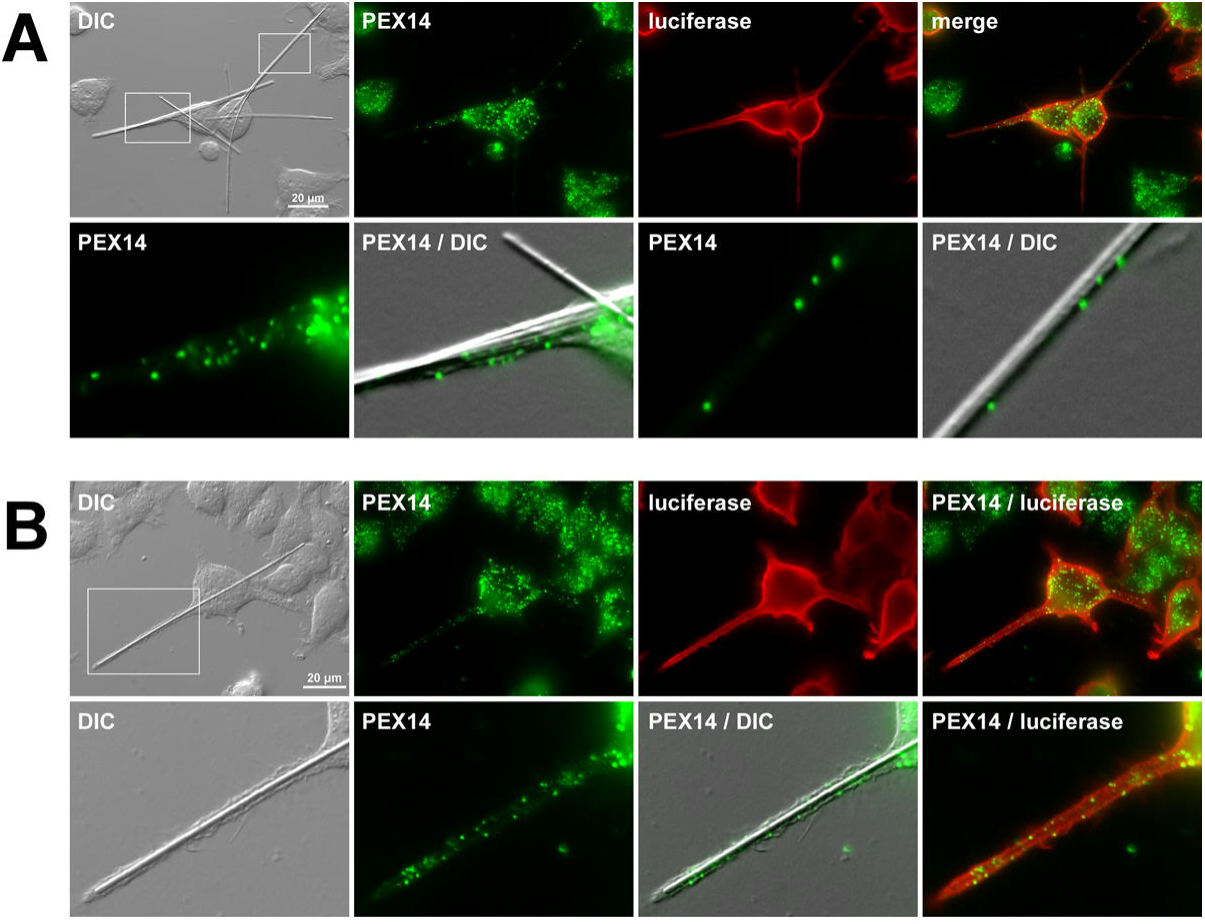
Needle-like inclusion bodies are not enclosed by peroxisome membranes. Transfected cells were seeded onto glass coverslips and moved from 37°C to 27°C culture condition 24 hours after transfection. On day-4 post-transfection, cells were fixed, permeabilized, and co-immunostained with rabbit anti-PEX14 polyclonal antibody (green) and goat anti-luciferase polyclonal antibody (red). Two representative image fields were shown. In the top row of A and B, the ‘areas of interest’ are marked in white boxes and are enlarged in the second row to show the relationship between the needle-shaped inclusion bodies and peroxisomes size, distribution, or morphology.

### 2.4. D-luciferin and its synthetic analog can inhibit intracellular luciferase crystallization

To find a condition or a factor that modulates luciferase crystallization in the 25°C–31°C temperature range, I examined the effect of D-luciferin on intracellular luciferase crystallogenesis. If a membrane-permeable substrate is added to the cell culture medium, the expectation is that the substrate will permeate into cells and interact with luciferase directly in the cytosol, or elsewhere. This, in turn, may influence the luciferase conformation, dynamics, stability, turnover, *etc*. thereby altering the crystal morphology or crystallizing propensity. In fact, when transfected HEK293 cells were continuously cultured in growth media containing 500 μM D-luciferin sodium salt or 500 μM CycLuc1 [10] (a synthetic luciferin) at 27°C, intracellular crystallization was completely prevented during the duration of cell culture for at least up to 5 days (Fig. 5A, second and third rows), while the DMSO (vehicle control) treated HEK293 cells readily induced the needle-shaped inclusion bodies (Fig. 5A, first row). Because D-luciferin and CycLuc1 treatment did not prevent the intracellular crystallization of unrelated proteins such as CRYGD (R37S mutant) and CLC, the blockade of luciferase crystallization was attributable to the interactions between luciferase and luciferin (data not shown). Because luciferase dynamically changes its conformation (including a 140° rotation of the small C-terminal domain [33, 34]) as part of its catalytic cycle in the presence of luciferin and co-factors ATP and Mg^2+^ [7, 35, 36], it is plausible that such processive conformational changes underscored the prevention of luciferase crystallization.

**Figure 5.**
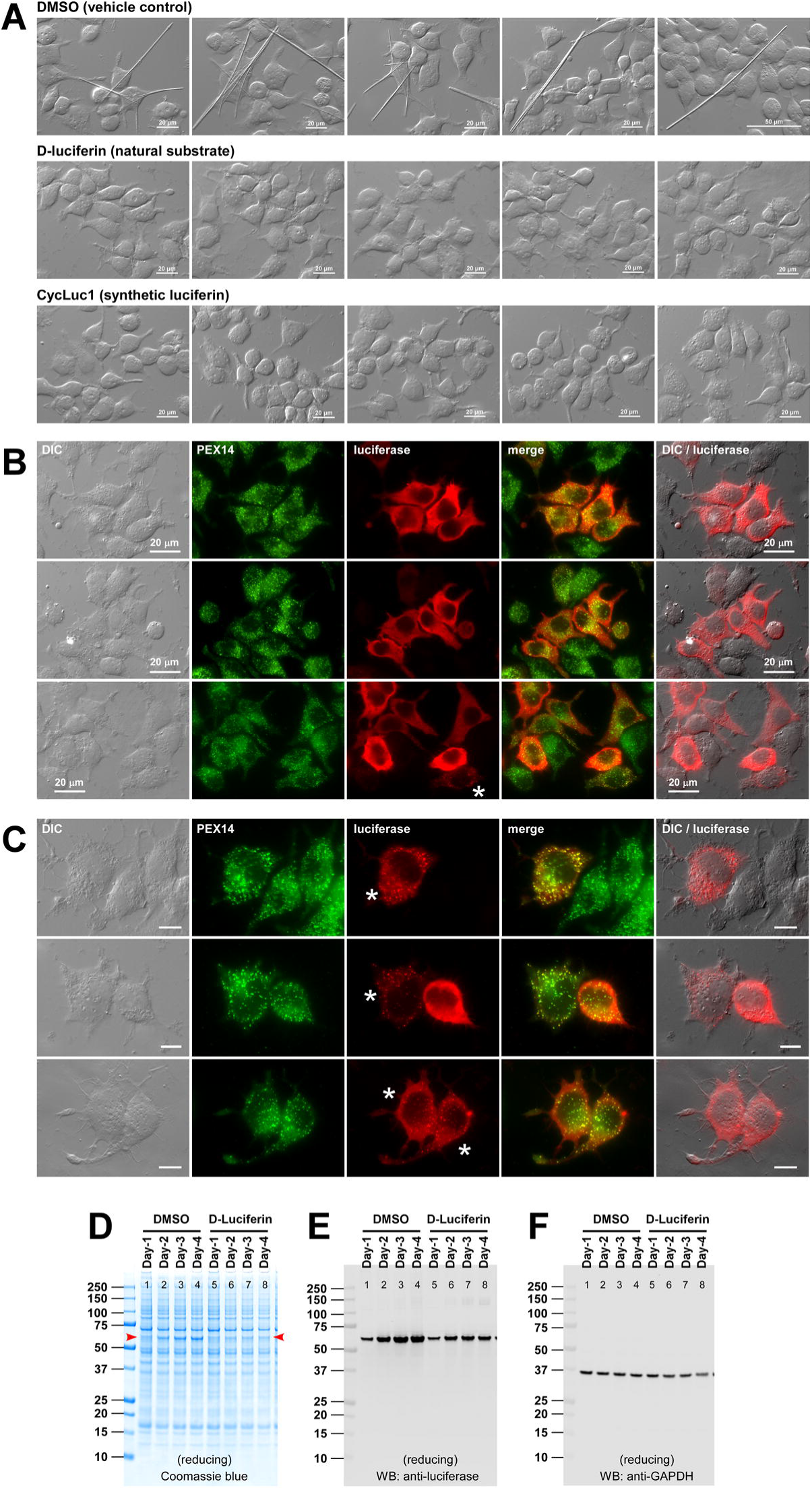
Intracellular luciferase crystallization is blocked by the presence of substrates. (A) Transfected cells were seeded onto glass coverslips at 24 hours post-transfection and were transferred from 37°C to 27°C culture condition in the presence of 0.5% (v/v) DMSO (first row), 150 μg/ml D-luciferin (second row), or 150 μg/ml CycLuc1 (third row). On day-4 post-transfection, cells were fixed, and the image fields were photographed using a DIC microscope. (B, C) On day-4 post-transfection, D-luciferin treated cells were fixed, permeabilized, and co-immunostained with rabbit anti-PEX14 (green) and goat anti-luciferase (red). Green and red image fields are digitally overlayed to create ‘merge’ views. DIC and red image fields were overlayed in the fifth column. Asterisks are placed next to the cells that show punctate luciferase staining. (D, E, F) Whole cell lysates were prepared from transfected cells from day-1 to day-4 post-transfection and analyzed by Coomassie staining (D) or by Western blotting (E, F). Cell lysate samples equivalent to 12,000-12,500 cells were loaded per lane. Cell lysates from DMSO vehicle treated cells were analyzed in lanes 1L4, while the cell lysates from D-luciferin treated cells are shown in lanes 5L8.

In the presence of a substrate that permeates into cells, overexpressed luciferase still largely maintained its cytosolic localization (Fig. 5B, red staining). Likewise, peroxisomes continued to show dispersed distribution as they normally would (Fig. 5B, green). Notably, however, a minor population of transfected cells started displaying punctate luciferase localization (Fig. 5B, bottom row, asterisks) when the cells were continuously grown in D-luciferin or CycLuc1 containing media. At higher magnification, such luciferase puncta clearly co-localized with the peroxisome marker PEX14 (Fig. 5C, asterisks). Although the mechanisms were unclear, the amount of cytosolic luciferase pool significantly diminished in those small numbers of cells, and this, in turn, helped unmask the peroxisome-localizing luciferase in those cells. The result concomitantly demonstrated that a fraction of luciferase was indeed correctly imported to peroxisomes in HEK293 cells although they were usually masked by the overwhelming amount of cytosolic luciferase pool.

To test if the overall amount of luciferase protein accumulation was impacted by the constant exposure to D-luciferin, steady-state luciferase protein level was examined in cell lysates prepared from DMSO and D-luciferin treated cells for 4 days at 31°C. In the vehicle-treated cell population, intracellular luciferase constantly increased over the period of 4 days (Fig. 5 DE, lanes 1–4). Given that the half-life of luciferase in mammalian cells was shown to be ∼3 hours [8], this result suggested that intracellular crystallization may be an effective way to accumulate cytosolic proteins by diverting them from protein turnover and sequestering into a more protected “storage” pathway. Conversely, in the presence of D-luciferin, the balance between new synthesis and degradation was at equilibrium as the luciferase level stayed nearly constant during the same period (Fig. 5 DE, lanes 5–8). Because the cell lysate sample loading was normalized by cell number, the endogenously expressed GAPDH level was comparable across the samples (Fig. 5F). There seem to be at least two possible scenarios as to how D-luciferin prevents intracellular luciferase crystallization. (i) Crystallization was prevented by D-luciferin because luciferase conformation changed dynamically during enzymatic cycles, and such dynamic movement hampered the intermolecular packing required for crystallization. (ii) In the presence of D-luciferin, the high turnover rate of luciferase was maintained and the luciferase did not have the chance to accumulate above a threshold concentration in the cytosol to initiate spontaneous crystallization.

### 2.5. Intracellular luciferase crystallization is prevented by luciferase inhibitors

Luciferase crystallization in HEK293 cells was also blocked when the transfected cells were cultured in the presence of luciferase inhibitors such as 2-(2-Hydroxyphenyl)benzothiazole (HPBT) [8] at 5 μg/ml or 20 μg/ml (Fig 6 ALC) and 2-phenylbenzothiazole (PBT) [8] at the same concentration range (data not shown). Because of their structural similarity to D-luciferin, these inhibitors complete with D-luciferin for the luciferin binding pocket and work as inhibitors of luciferase in biochemical assays in test tubes [8, 9]. However, HPBT and related compounds are known to extend the luciferase’s intracellular half-life inside the mammalian cells through a mechanism called “inhibitor-based stabilization” and paradoxically enhance bioluminescence in cell-based assays [5, 8, 37]. In agreement with such a mode of action, intracellular luciferase levels continued increasing over the period of 5 days in HPBT-treated cells (Fig. 6 DE, lanes 5– 12), and the increase was indistinguishable from the DMSO-treated cells that fostered crystal growth (Fig. 6 DE, lanes 1–4). Again, the cell lysate sample loading was normalized by cell number, and the endogenously expressed GAPDH level was comparable across the samples (Fig. 6E, bottom panel). The underlying mechanics of how HPBT and PBT blocked intracellular luciferase crystallization were, therefore, different from those of D-luciferin and CycLuc1. It is likely that HPBT binds to the luciferin binding pocket and locks luciferase into a stable yet crystallization-incompetent conformation.

**Figure 6.**
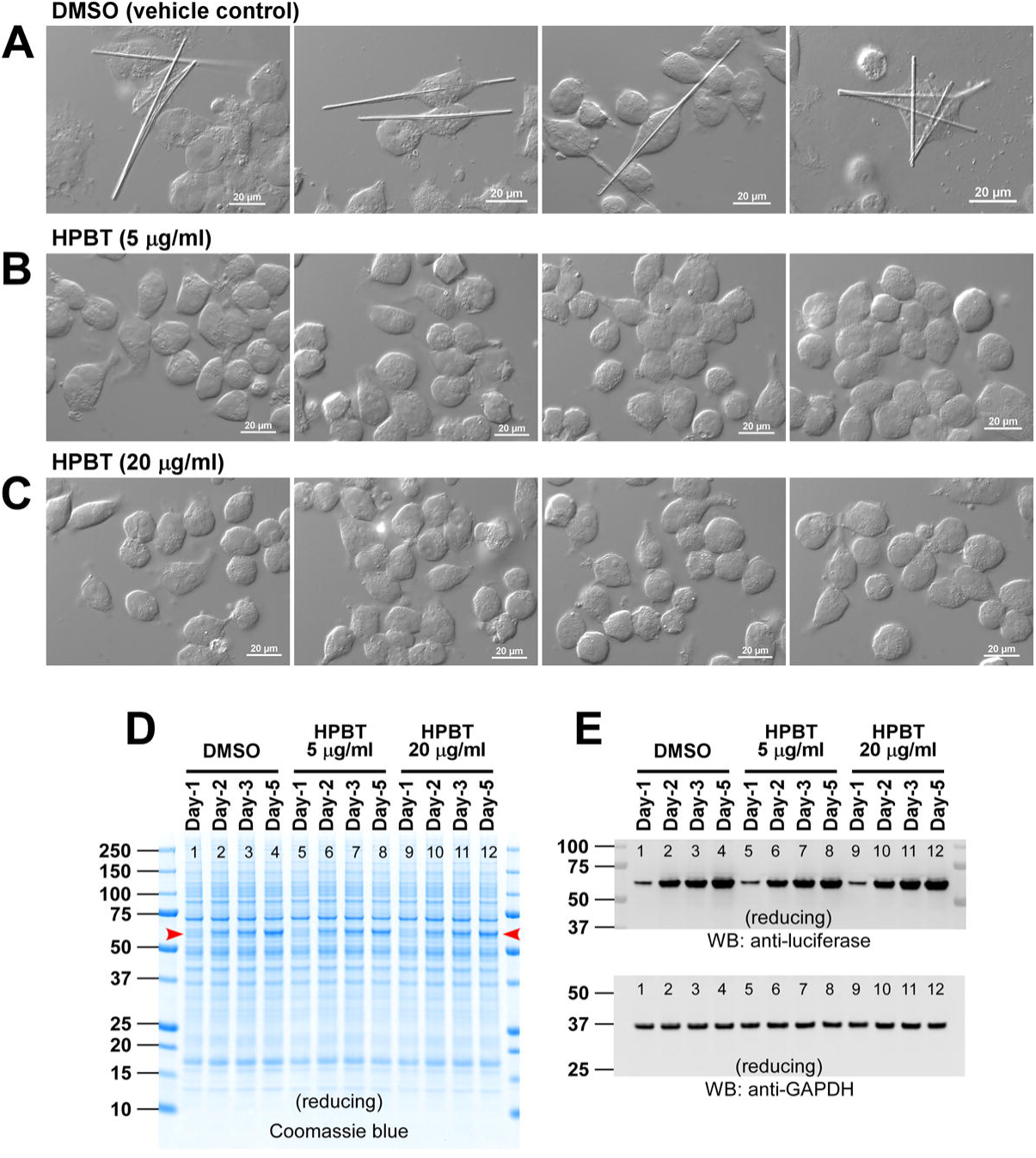
Intracellular luciferase crystallization is blocked by hydroxyphenyl benzothiazole (HPBT). (A, B, C) Transfected cells were seeded onto glass coverslips at 24 hours post-transfection and transferred from 37°C to 27°C culture condition in the presence of (A) 0.5% (v/v) DMSO, (B) 5 μg/ml HPBT, or (C) 20 μg/ml HPBT. On day-4 post-transfection, cells were fixed, and images were photographed using a DIC microscope. (D, E) Whole cell lysates were prepared from transfected cells from day-1 to day-5 post-transfection and analyzed by Coomassie staining (D) and by Western blotting (E). Cell lysates corresponding to 12,000-12,500 cells are loaded and analyzed per lane. Cell lysates from DMSO treated cells were analyzed in lanes 1L4, while cell lysates from 5 μg/ml HPBT and 20 μg/ml HPBT were shown in lanes 5L8 and lanes 9L12, respectively. Membranes were probed by anti-luciferase antibody (top row) or anti-GAPDH (bottom row).

## 3. Discussion

This study uncovered unexpected characteristics of firefly luciferase that temperature-dependently phase separates into needle-shaped crystal-like inclusion bodies in the cytosol of transfected HEK293 cells. Because firefly luciferase is imported into the lumen of peroxisomes post-translationally after completing its biosynthesis in the cytosol [38], when the rate of protein synthesis exceeds the capacity of the PTS1 pathway, luciferase progressively accumulates in the cytosol. Although the overexpressed firefly luciferase accumulated in the cytosol of transfected HEK293 cells to a similar level both at normal cell culture temperature of 37°C and hypothermic temperatures (25°CL31°C), luciferase behaved differently depending on the cell culture temperatures. While luciferase aggregated into a large aggresome-like protein mass in the cytosol at 37°C in roughly half of the transfected cell population, luciferase spontaneously produced needle-shaped crystal-like inclusion bodies at hypothermic temperatures. The high rate of luciferase aggregation at 37°C suggested that luciferase concentration in the cytosol was already near the upper limit of colloidal stability and solubility. Therefore, insufficient luciferase concentration was not the likely reason why luciferase did not crystallize at 37°C. Rather, the results indicated that the hypothermic temperatures somehow influenced the conformation or molecular dynamics of firefly luciferase in such ways as to favor the spontaneous formation of crystal-like inclusion bodies. Intracellular protein crystallization has been believed to occur very rarely in nature [39], but the list of such examples is growing [40, 41]. Current research inadvertently identified firefly luciferase as an excellent model protein for studying the intracellular liquid-to-solid phase transition process triggered by a simple cell culture temperature alteration.

Another unexpected finding of firefly luciferase phase behavior was the prevention of intracellular crystal-like inclusion formation by the substrate D-luciferin and CycLuc1. In addition, so-called “stabilizing inhibitors” belonging to benzothiazoles prevented the needle-like inclusion body formation. Although D-luciferin and luciferase inhibitors are structural analogs, their mode of action in blocking the intracellular luciferase crystallization was different. In the presence of D-luciferin, the amount of intracellular luciferase remained nearly constant throughout the testing period up to day-5 post-transfection. This suggested that luciferase’s new synthesis and turnover were balanced. By contrast, in the presence of benzothiazole class inhibitors, intracellular luciferase levels steadily increased over time during the same testing period as expected from their luciferase-stabilizing effects in cell-based systems [8]. Therefore, D-luciferin appeared to prevent the luciferase crystallization by suppressing the cytosolic accumulation of luciferase above threshold and by inducing dynamic conformation changes, whereas benzothiazoles stabilized the luciferase conformation into a crystallization-incompetent state that resisted efficient protein turnover. This study suggests that, under substrate insufficiency, the excess firefly luciferase phase separates into a crystal-like state that can modulate protein turnover rate and intracellular soluble enzyme availability.

Immunofluorescent staining clearly indicated that the crystal-like inclusion bodies were not associated with or encapsulated by membrane-bound organelles. Similarly, the inclusion bodies did not show an association with proteinaceous polymeric structures such as microtubule networks. The needle-shaped inclusion bodies were, therefore, likely produced from the cytosolic luciferase pool. In many cases, the length of needles exceeded 100 μm in length, and the mechanical tension produced by the elongating needles extended the cell morphology longitudinally. In other words, the inclusion body was contained within the boundary of the plasma membrane, and its integrity was maintained despite the mechanical stretch stress from the elongating inclusion bodies. Previously, Schönherr et al. [24] reported that similar intracellular luciferase crystals produced in Sf9 insect cells were surrounded by the peroxisome membranes shown by PEX26-mCherry protein. In the present study, peroxisomes were visualized by staining an endogenous peroxisome protein PEX14, yet the association between peroxisomes and the luciferase needles was not observed. In fact, individual peroxisomes themselves were by no means large enough to enclose the needle-like inclusion bodies. If peroxisomes were to encapsulate the needles, individual peroxisomes must homotypically fuse to generate a large membranous structure, but such a membrane fusion event was not evident in HEK293 cells.

It is interesting that the hypothermic cell culture settings coincide with the temperature range of firefly habitats in temperate regions. This brings up the following question: Does firefly luciferase crystallize into needle-shaped inclusion bodies in the light-producing photocytes of the firefly lantern? From the physiological and evolutional standpoints, it is easy to envision that such an event needs to be suppressed because, otherwise, luciferase crystals would not only physically damage the photocytes and the lantern organ but also diminish the mating opportunity or even threaten the life of the firefly. In this sense, some suppression mechanisms must be evolutionarily built into the primary sequence or the cellular environment in which luciferase is expressed [39]. Then, how is luciferase crystallization suppressed in the firefly photocytes? The present study suggested a few of such potential failproof mechanisms. Firstly, to ensure that synthesized luciferase does not over-accumulate in photocytes, the expression level of luciferase must be carefully controlled *in vivo*. If this step functions properly, luciferase does not crystallize in photocytes, although a firefly’s temperate habitat may be favorable for luciferase protein to crystallize. Secondly, the firefly lantern photocytes are densely filled with peroxisomes [13, 42, 43], so there should be sufficient intrinsic capacity to transport luciferase into the peroxisome lumen via the PTS1 pathway to prevent the accumulation in the cytosol. However, given the pH of the cytosol (∼7.2) and peroxisomal lumen (∼6.9L7.0) is similar [44, 45] if luciferase ends up accumulating in peroxisome to a high enough concentration, it may still crystallize at a suitable temperature range. Therefore, an additional layer of preventive measures must be in place. What can that be? This study demonstrated that intracellular crystallization of firefly luciferase was effectively inhibited by the presence of substrate D-luciferin. Because the firefly lantern photocytes synthesize and contain both D-luciferin substrate and luciferase enzyme in peroxisomes [12, 13, 46], it is highly likely that intracellular luciferase crystallization is suppressed in the light-emitting photocytes by conformation dynamics that disfavor crystallization and/or by ensuring a steady enzyme turnover to discourage accumulation. Luciferase crystallization can thus be steadfastly suppressed in photocytes by virtue of a multi-tiered approach by regulating protein synthesis, peroxisome import, conformation, and turnover rate.

Interestingly, however, protein crystals are very prevalent in bioluminescent organisms. For instance, although the protein identities have not been investigated in detail yet, some species of fireflies (e.g., *Photinus macdermotti* [13] and *Photinus Greeni* [47]) frequently house small protein crystals inside peroxisomes in their lantern photocytes. In single-celled bioluminescent marine dinoflagellate *Gonyaulax polyedra*, luminescent blue light is produced in birhombohedral crystalline luminous particles named “scintillons” [48], which are composed of 135 kDa luciferase and 75 kDa luciferin-binding protein [49]. Furthermore, in the Cephalopod named Japanese “firefly squid” *Watasenia scintillans*, the bright blue bioluminescent signal is generated in the three small light organs called photophores located at the tips of the pair of fourth arms [50, 51]. These light-producing photophores are filled with numerous rod-shaped crystalline bodies composed of three related luciferase-like proteins, wsluc1–3 [52]. Likewise, in the ocular light organ of “lyre cranch squid” *Bathothauma lyromma*, photophores are filled with a mass of protein crystals [53]. Whether these protein crystals are mere co-incidence among different bioluminescent organisms is currently unknown. However, it is attractive to speculate that a group of proteins involved in bioluminescence are prone to crystallization as if it is a shared theme for the light organ evolution in terrestrial and marine bioluminescent organisms.

Why luciferase crystallization in mammalian cells had been unrecognized despite the widespread use of firefly luciferase in numerous biomedical and life science applications? While there is a possibility that nobody took the time to report it even though this event has been known widely, there are a few potential explanations for why such a phenomenon has escaped researchers’ attention. For instance, (a) in most applications, engineered mammalian cells that express luciferase reporter are cultured and tested at 37°C or grafted to animal models *in vivo*. At this normal temperature range of mammalian cells and animals, luciferase crystallization is effectively suppressed. (b) In cell-based reporter gene assays, luciferase expression is often driven by a responsive element active only under certain signaling settings. As a result, unlike the case of strong constitutive viral promoters/enhancers, luciferase protein expression would not reach a high enough level to induce intracellular crystallization. (c) The vast majority of cell-based luciferase reporter assay applications depend on a cell lysis step. (d) Few scientists in the past had good reasons to culture luciferase-expressing mammalian cells at a hypothermic temperature range from 25°C to 31°C for morphological studies.

Mild hypothermic cell culture conditions, especially at around 31°C, are widely used in the biotechnology industry when monoclonal antibodies and other secretory proteins of therapeutic values are produced from stably transfected CHO cells [54, 55]. Such conditions have proven effective in prolonging cell health and slowing down cell doubling while diverting energy to protein expression to enhance specific productivity and tune desired quality attributes [55, 56]. Similarly, anyone who has hands-on experience in E.coli recombinant protein expression systems would attest that when your proteins are insolubly aggregated at 37°C induction temperature, one can employ lower temperatures as low as 18°C to recover the protein of interest in the soluble fraction [57]. The example of firefly luciferase intracellular crystallization only at hypothermic culture conditions and led to abundant intracellular storage suggests that there is much more room to operate for mammalian cell growth conditions when optimizing high-value recombinant protein production. Historically, it is well known that mammalian cell’s secretory activity stops if the cell culture temperature goes below 20°C [58]. In this sense, it is reasonable to go beyond normal practice and optimize cell culture temperatures between 20°C and 37°C regardless of secretory or intracellular protein productions. Such a simple yet potentially impactful cell culture approach may open new possibilities for mammalian cells as an even more versatile system to produce broad families of recombinant proteins.

## 4. Materials and Methods

### 4.1. Detection antibodies and reagents

Goat polyclonal anti-luciferase was purchased from Promega (cat. G7451) and used for immunostaining and immunoblotting. Rabbit anti-luciferase monoclonal antibody [EPR17789] was from abcam (cat. ab185923) and used for immunostaining. Rabbit polyclonal anti-PEX14 was obtained from ProteinTech (cat. 10594-1-AP). Rabbit anti-calnexin (cat. C4731) and rabbit anti-TGN46 (cat. T7576) were from Sigma-Aldrich. Mouse anti-p230 was from BD Bioscience (cat. 611280). Rabbit anti-giantin was from Covance (cat. PRB-114P). Mouse anti-LAMP1 (clone H4A3, cat. sc-20011) and rabbit anti-Tom20 (FL-145, cat. sc-11415) were from Santa Cruz Biotechnology. Mouse anti-β-tubulin (E7) was from the Hybridoma bank. Mouse anti-Nup153 (clone QE3, cat. 906201) was from Biolegend. Mouse anti-GAPDH (clone 6C5) was from Chemicon (cat. MAB374). D-Luciferin sodium salt (cat. L6882), CycLuc1 (cat. 5.30650), 2-(2-Hydroxyphenyl)benzothiazole (cat. 632589), and other chemicals were obtained from Sigma-Aldrich unless specifically mentioned.

### 4.2. Expression constructs

The nucleotide sequence encoding *Photinus pyralis* (North American firefly) luciferase was the same as GenBank AB644228. The recombinant DNA sequence was verified by the Sanger method and subcloned into a pTT^®^5 mammalian expression vector licensed from the National Research Council of Canada.

### 4.3. Cell culture and transfection

HEK293-6E cell line (herein HEK293) was obtained from the National Research Council of Canada. HEK293 cells were suspension cultured in a humidified CO2 Reach-In incubator (37 °C, 5% CO2) using FreeStyle™ 293 Expression Medium (Thermo Fisher Scientific). The cells were grown in vented cap Corning^®^ Erlenmeyer flasks placed on Innova 2100 shaker platforms (New Brunswick Scientific) rotating at 130L135 rpm. The expression construct was transfected into HEK293 cells using the protocol described in detail previously [59]. Difco yeastolate cell culture supplement (BD Biosciences) was added to the suspension cell culture at 24 hr post-transfection. Depending on the nature of temperature shift experiments, the flasks of transfected cells were moved directly to a shaker platform located in an insect cell culture incubator (27°C, ambient CO2) or a Reach-In CO2 incubator at 31°C, 5% CO2. For imaging studies, transfected cells were seeded onto glass coverslips at 24 hours post-transfection and statically cultured in cooling CO2 incubators (ESCO CelCulture^®^) adjusted at 37°C, 31°C, 27°C, or 25°C in the presence or absence of designated test compounds for up to day-5 post-transfection.

### 4.4. Microscopy

To perform imaging experiments, transfected cells in suspension format were seeded onto poly-D-lysine coated glass coverslips placed in 6-well plates at 24 hours post-transfection. Cells were then statically cultured for up to day-6 post-transfection in CO2 incubators at 37°C, 31°C, 27°C or 25°C in the presence or absence of test compounds. Cells were fixed in 100 mM sodium phosphate buffer (pH 7.2) containing 4% paraformaldehyde for 30 min at room temperature. After three washing/quenching steps in PBS containing 0.1 M glycine, fixed cells were either directly used for DIC or phase contrast microscopy or processed for indirect immunofluorescent microscopy. To carry out immunostaining, cells were permeabilized in PBS containing 0.4% saponin, 1% BSA, and 5% fish gelatin for 15 min, followed by an incubation with primary antibodies diluted in the same permeabilization buffer for 60 min. After three washes in the permeabilization buffer, the cells were probed with secondary antibodies for 60 min in the permeabilization buffer. Coverslips were mounted onto slide-glass using Vectashield mounting media (Vector Laboratories) and cured overnight at 4°C. The slides were analyzed on a Nikon Eclipse 80i microscope with a 60× or 100× CFI Plan Apochromat oil objective lens and Chroma FITC-HYQ or Texas Red-HYQ filter. DIC and fluorescent images were acquired using a Cool SNAP HQ2 CCD camera (Photometrics) and Nikon Elements BR imaging software. Zeiss AXIO Imager.A1, equipped with an AxioCam CCD camera, was used for phase contrast microscopy.

### 4.5. SDS-PAGE and Western blotting

At a designated time point after transfection, an aliquot of suspension cell culture was removed from shake flasks, and the cells were harvested with a centrifugation step at 1,200 g for 6 min. Cell pellets were directly lysed in a lithium dodecyl sulfate sample buffer (Thermo Fisher Scientific) containing 5% (v/v) β-mercaptoethanol and heat treated for 5 min at 75°C. To normalize the sample loading, whole cell lysates corresponding to 12,000–12,500 cells were analyzed per lane. SDS-PAGE was performed using NuPAGE 4–12% Bis-Tris gradient gel and a compatible MES SDS buffer system (both from Thermo Fisher Scientific). Resolved proteins were electrotransferred to a nitrocellulose membrane, blocked with fluorescent Western blocking buffer (Rockland Immunochemicals), and probed with designated primary antibodies. After three washes in PBS containing 0.05% (v/v) Tween-20, the nitrocellulose membranes were probed with Alexa Fluor-680 conjugated secondary antibody (Thermo Fisher Scientific). The images of fluorescent Western blot were acquired using an Odyssey^®^ infrared imaging system (LI-COR Biosciences).

## 6. Acknowledgements

The author acknowledges Elham Ettehadieh for making an expression vector encoding firefly luciferase. I thank Jason Lam for the valuable discussion on peroxisome protein import. HH is personally grateful to Yoko Azumi for her support and encouragement.

